# Intergenerational Conditioning via Intermittent Parental Hypoxia Confers Stroke Resilience in Offspring

**DOI:** 10.1101/2025.07.02.662697

**Authors:** Ahmet B. Caglayan, Mustafa C. Beker, Favour Felix-Ilemhenbhio, Serdar Altunay, Hayriye E. Yelkenci, Aysun Caglayan, Enes Dogan, Nilay Ates, Ok-Nam Bae, David J. Burrows, Ali Ali, Milena De Felice, Ertugrul Kilic, Arshad Majid

## Abstract

**Background and Aims:** Intergenerational disease transmission, where parental exposures or experiences influence disease susceptibility in offspring, may represent a crucial layer of stroke risk that extends beyond genetics alone. Environmental conditioning, such as intermittent sub-lethal hypoxia, can induce adaptive protective stress responses in the brain. However, whether such parental conditioning enhances offspring resilience to cerebral ischaemia remains unclear. This study investigates whether intermittent hypoxia in parents acts as an intergenerational conditioning stimulus, conferring resilience to ischaemic stroke in offspring, and explores associated molecular mechanisms.

**Methods:** Male and female Balb/C mice (F0) were exposed to intermittent hypoxia (8% O_2_, 2 hours every other day, 16 cycles) prior to mating. To confirm that intermittent hypoxia induced neuroprotection in the parental generation, a separate cohort of F0 mice underwent transient middle cerebral artery occlusion (tMCAO). Offspring (F1) were generated from hypoxia-exposed F0 breeders and divided into four groups: biparental hypoxia, paternal hypoxia, maternal hypoxia, and normoxic controls. Adult F1 offspring also underwent tMCAO to model ischaemic stroke. Infarct volume and brain swelling were assessed 48 hours post-ischaemia. In a subgroup of F1 offspring, tandem mass tag (TMT)-based proteomic analysis of injured brain tissue was performed post-stroke to identify molecular pathways associated with neuroprotection.

**Results:** Parental intermittent hypoxia significantly reduced infarct size and swelling in F0 mice. These protective effects were inherited by F1 offspring, with biparental exposure producing the greatest reduction in infarct volume, followed by maternal-only and paternal-only groups, and exhibiting sex-specific differences. Proteomic profiling revealed distinct treatment and lineage clusters. Key pathways implicated in offspring neuroprotection included metabolic regulation, immune signalling, cytoskeletal organisation, and cell survival, notably involving PI3K-Akt and EGFR pathways.

**Conclusions:** Intermittent hypoxia in parents acts as an intergenerational conditioning stimulus, conferring offspring resilience to ischaemic stroke. This neuroprotective phenotype is supported by coordinated molecular adaptations in key pathways involved in survival and stress response. These findings highlight the potential for parental environmental conditioning to shape stroke outcomes in offspring, opening new avenues for therapeutic exploration.

## 1. Introduction

Ischaemic stroke is a leading cause of adult disability worldwide, and despite advances in acute management, outcomes remain highly variable. While genetic factors undoubtedly contribute to stroke susceptibility, a growing body of evidence suggests that intergenerational disease transmission, in which parental exposures or experiences influence offspring vulnerability, plays an important role in determining stroke risk and severity (1).

One potential mechanism by which parents might influence offspring stroke outcomes is via environmental conditioning (2). Intermittent sub-lethal hypoxia is a well-established preconditioning stimulus that can induce neuroprotective adaptations in the brain, largely through activation of stress response pathways, improved metabolic efficiency, and modulation of cell survival mechanisms (2, 3). However, whether parental exposure to intermittent hypoxia can confer intergenerational resilience to ischaemic injury in offspring has not been fully explored.

Here, we investigate whether parental intermittent hypoxia acts as an intergenerational conditioning stimulus to enhance offspring resilience to ischaemic stroke. Using a well-established mouse model of transient middle cerebral artery occlusion (tMCAO) (4), we assess infarct volume and brain swelling in offspring from different parental hypoxia exposures. We further explore the underlying molecular mechanisms through proteomic analysis to identify key pathways associated with neuroprotection.

## 2. Materials and Methods

### Animals and Experimental Procedures

All procedures were approved by the Ethics Committee of Istanbul Medipol University (2020/16) and conducted in accordance with the Declaration of Helsinki and relevant institutional guidelines. Male and females Balb/C mice (8–10 weeks old) were housed under a 12:12-hour light–dark cycle with ad libitum access to food and water.

### Hypoxia Induction and Breeding

Intermittent hypoxia was induced using a plexiglass chamber equipped with controlled heating and exhaust, supplied with 8% O_2_ balanced with N_2_ (5). Mice were exposed to hypoxia for 2 hours every other day over 16 cycles. Normoxic controls were exposed to 21% O_2_ under the same conditions. After hypoxia, one cohort of F0 mice (aged 12 weeks) was used for immediate stroke injury evaluation, while another cohort was bred to generate F1 offspring (**Figure 1A**).

**Figure 1.**
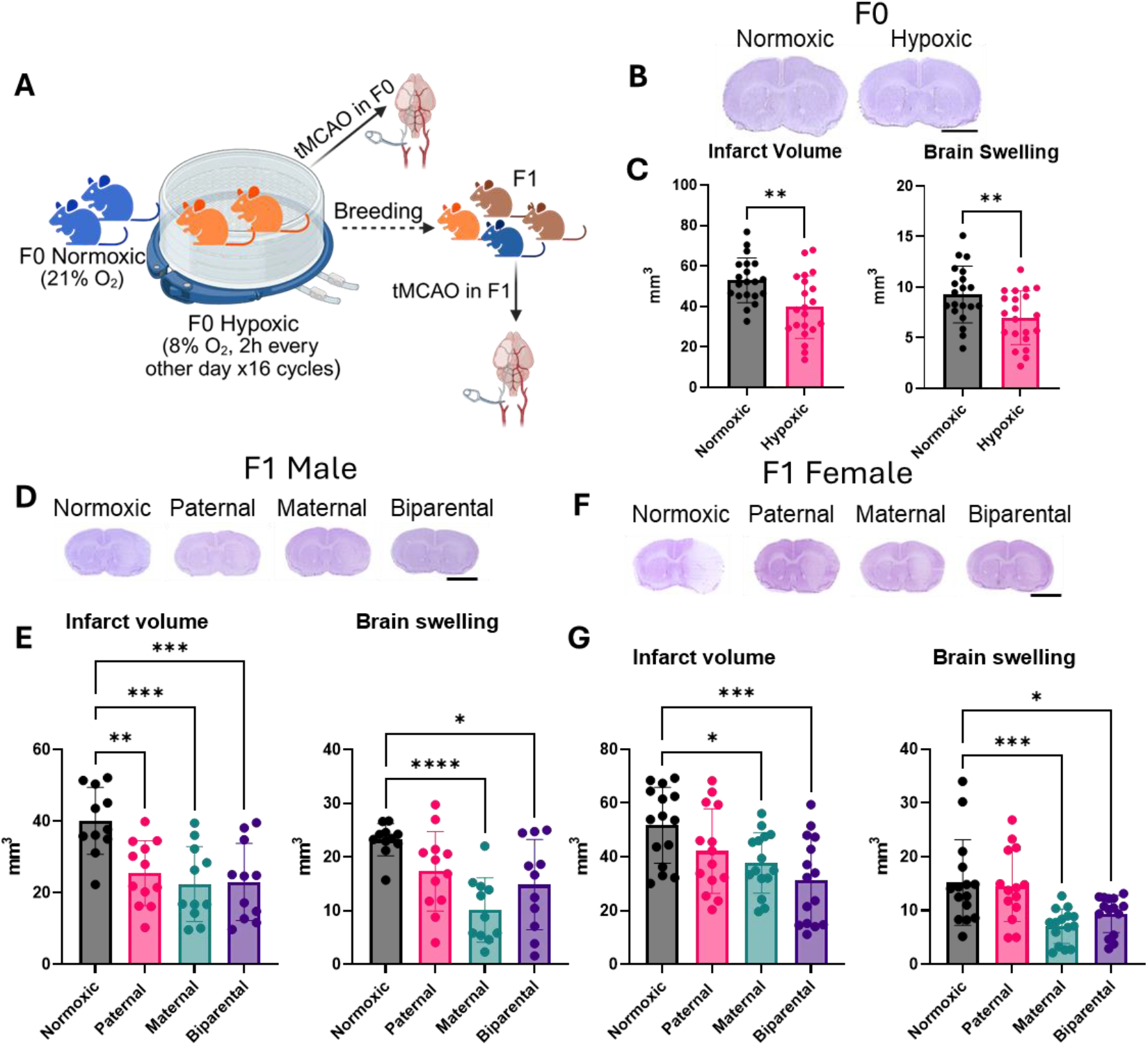
Parental intermittent hypoxia reduces infarct volume and brain swelling following ischaemic stroke in F0 and F1 mice. **A)** Schematic of the in vivo mouse model. F0 mice were exposed to intermittent hypoxia prior to tMCAO. A subset was bred to generate F1 offspring, which also underwent tMCAO to assess intergenerational effects. Image created with BioRender. **B**) Representative coronal brain sections stained with cresyl violet showing infarcted regions in normoxic and hypoxia-exposed F0 mice (scale bar = 2.5 mm). **C)** Quantification of infarct volume and brain swelling in F0 mice (N=20-21; equal male and female distribution). Data are shown as mean ± SEM. Statistical analysis: unpaired *t*-test. **D)** Representative cresyl violet-stained brain sections from F1 male offspring of normoxic and hypoxia-exposed breeders (scale bar = 2.5 mm). **E)** Quantification of infarct volume and brain swelling in F1 male offspring (N=11-12). Analyses are one-way ANOVA with Dunnett’s multiple comparisons test. **F)** Representative cresyl violet-stained brain sections from F1 female offspring (scale bar = 2.5 mm). **G)** Quantification of infarct volume and brain swelling in F1 female offspring (N=14-15).

### Transient Middle Cerebral Artery Occlusion (tMCAO)

Transient MCAO was induced using the intraluminal filament technique (4). Mice were anaesthetised with 1.5% isoflurane (30% O_2_, remainder N_2_O), and body temperature was maintained at 36.5–37°C. Cerebral blood flow was monitored using laser Doppler flowmetry to confirm occlusion. The left common and external carotid arteries were ligated, and a filament was introduced through the common carotid artery to occlude the MCA. After 60 minutes, reperfusion was initiated by filament removal, and cerebral blood flow was monitored for an additional 20 minutes. Both a cohort of hypoxia-exposed F0 mice (aged 12 weeks) and adult F1 offspring (aged 8-10 weeks) underwent tMCAO.

### Infarct Volume and Brain Swelling Assessment

At 2 days post-ischaemia, brains were sectioned coronally at 2 mm intervals and stained with cresyl violet. Infarct areas were calculated by subtracting the non-infarcted ipsilateral hemisphere area from the contralateral hemisphere using ImageJ. Infarct volume was integrated across sections. Brain swelling was assessed by measuring the volume difference between hemispheres (6, 7).

### Proteomics

Injured cortex samples from F1 mice at 55 days post-stroke were collected, homogenised, and lysed for protein extraction (8). Proteins (50 μg) were processed using filter-aided sample preparation (FASP), alkylated, and digested overnight with trypsin. LC-MS/MS analysis was performed on a UPLC system coupled to a SYNAPT G2-Si mass spectrometer. Data processing was conducted with Progenesis QI, requiring ≥2 unique peptides for protein identification. Downstream data analysis and visualisation were conducted in R (v4.5.0): heatmaps (pheatmap v1.0.12), general plots (ggplot2 v3.5.1), volcano plots (EnhancedVolcano v1.20.0), and Venn diagrams (ggVennDiagram v1.5.2). KEGG pathway enrichment analysis was performed using clusterProfiler (v4.17.0) and org.Mm.eg.db (v3.21.0).

### Statistical Analysis

Data were analysed using t-tests or one-way ANOVA with LSD post hoc tests, as appropriate, in GraphPad Prism (v10.4.2) and R. Results are reported as mean ± SEM, with p < 0.05 considered statistically significant.

## 3. Results

### Hypoxic preconditioning in F0 mice reduces ischaemic brain injury

To determine the effect of intermittent hypoxia on stroke outcomes, we compared infarct volumes and brain swelling in F0 mice subjected to tMCAO. As previously demonstrated by us and others, intermittent hypoxia significantly reduced infarct volume (25.01% reduction; *p* = 0.0032) and brain swelling (24.86% reduction; *p* = 0.0099) compared to normoxic controls **(Figure 1B-C)**.

Breeding from hypoxia-exposed parents produced four F1 groups: paternal-only, maternal-only, biparental hypoxia, and normoxic controls. Following tMCAO in adult F1 offspring, all hypoxic groups exhibited reduced infarct volume and brain swelling, with sex-specific differences **(Figure 1D-G)**.

In males (**Figure 1D-E**), infarct volume was significantly reduced in the paternal (37.5% reduction; *p* = 0.0030), maternal (45% reduction; *p* = 0.0005), and biparental (42.5% reduction; *p* = 0.0007) groups compared to normoxic controls. However, only the maternal (56.52% reduction; *p* < 0.0001) and biparental (34.78% reduction; *p* = 0.0125) groups showed significantly reduced brain swelling; paternal hypoxia showed a non-significant trend (26.10% reduction; *p* = 0.0929).

In females (**Figure 1F-G**), infarct volume was significantly reduced in the maternal (26.92% reduction; *p* = 0.0272) and biparental (40.38% reduction; *p* = 0.0007) groups, with a non-significant trend in the paternal group (19.23% reduction; *p* = 0.1883). Brain swelling showed a similar pattern, with significant reductions in the maternal (53.33% reduction; *p* = 0.0008) and biparental (40.0% reduction; *p* = 0.0185) groups, but no reduction in the paternal group compared to normoxic controls (Figure 1C-F).

### Proteomic insights into intergenerational neuroprotection

Building on the observed neuroprotection in both F0 and F1 mice, we performed proteomic analysis to explore underlying molecular mechanisms. For F0 mice, we analysed both male and female brains. For F1 offspring, we focused on females due to the more consistent protective phenotype observed in the maternal and biparental hypoxia groups across both infarct volume and brain swelling. This targeted approach aimed to maximise the likelihood of detecting relevant proteomic changes associated with intergenerational neuroprotection.

Unsupervised hierarchical clustering of the proteomic data revealed clear segregation by treatment and lineage (Figure 2A). F0 and F1 normoxic female samples clustered together, indicating consistency across generations in the absence of hypoxic preconditioning. In contrast, F1 hypoxic groups (maternal, paternal, and biparental) formed a distinct cluster, suggesting coordinated protein expression changes linked to ancestral hypoxia exposure. Interestingly, F0 hypoxic and normoxic males clustered together, indicating limited proteomic alterations, while F0 hypoxic females diverged from normoxic controls, highlighting a sex-specific response to intermittent hypoxia.

**Figure 2.**
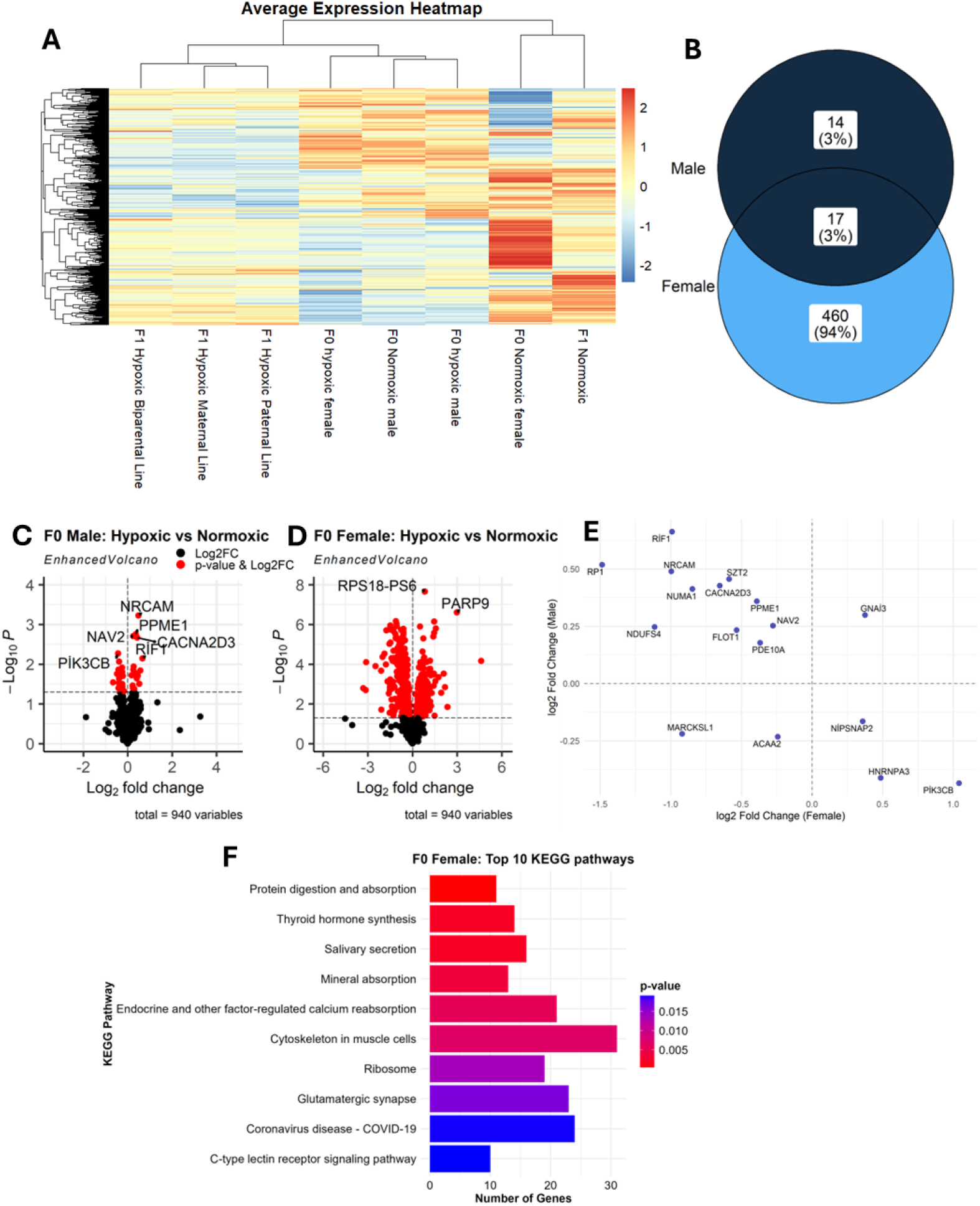
Proteomic analysis of hypoxia-induced changes in F0 mice brain cortex. **A)** Heatmap displaying scaled average protein expression across sample groups (N=6-8). Each row represents a protein, and each column represents a sample group. Expression values were averaged per group and scaled by protein (row-wise z-score). **B)** Venn diagram illustrating the number of differentially expressed (DE) proteins common to F0 male and female hypoxic mice compared to normoxic controls. **C-D)** Volcano plot showing DE proteins in hypoxic vs normoxic F0 male **(C)** and female **(D)** mice. Differential expression was defined by p-value < 0.05. **E)** Scatter plot showing the 17 DE proteins shared between hypoxic F0 male and female mice. **F)** KEGG pathway enrichment analysis showing the top 10 significantly enriched pathways for proteins differentially expressed in hypoxic vs normoxic F0 female mice.

### Differential proteomic responses in F0 hypoxic mice

Consistent with the hierarchical clustering, differential expression analysis revealed that hypoxia exposure altered the expression of 31 proteins in F0 males and 477 proteins in F0 females compared to their respective normoxic controls (*p* < 0.05; Figure 2B–E). Seventeen proteins were commonly differentially expressed in both sexes, suggesting a shared molecular response (Figure 2B). Among these, *Gnai3* was upregulated and *Marcksl1* was downregulated in both hypoxic males and females. Notably, *Gnai3* has been implicated in cerebral ischaemic injury and identified as a target of Astragaloside IV in a rat stroke model (9), highlighting its potential relevance to neuroprotection. *Marcksl1* has been identified as a differentially expressed gene in the mouse corpus callosum of ageing mice (10), suggesting a potential role in age-related neurological disorders.

KEGG pathway enrichment analysis of differentially expressed proteins (DEPs) in F0 females revealed significant enrichment in pathways related to protein digestion and absorption, thyroid hormone synthesis, mineral absorption, endocrine signalling, and cytoskeletal regulation (Figure 2F). Although these pathways are not directly linked to ischaemic injury, they may reflect broader systemic or metabolic adaptations to intermittent hypoxia. We focused on F0 females because they showed the most distinct clustering from normoxic controls and the highest number of DEPs, suggesting a more pronounced proteomic response to hypoxic preconditioning.

### Differential proteomic responses in F1 hypoxic mice

Differential expression analysis revealed that parental hypoxia exposure altered the expression of 44 proteins in the paternal group, 33 proteins in the maternal group, and 58 proteins in the biparental group compared to F1 normoxic controls (Figure 3A-E). Six proteins were commonly downregulated across all three groups, suggesting a shared molecular response (Figure 2E). These included *Akr1a1, Atp12a, Csde1, Gnaz, Impa1*, and *Srgap3*.

**Figure 3.**
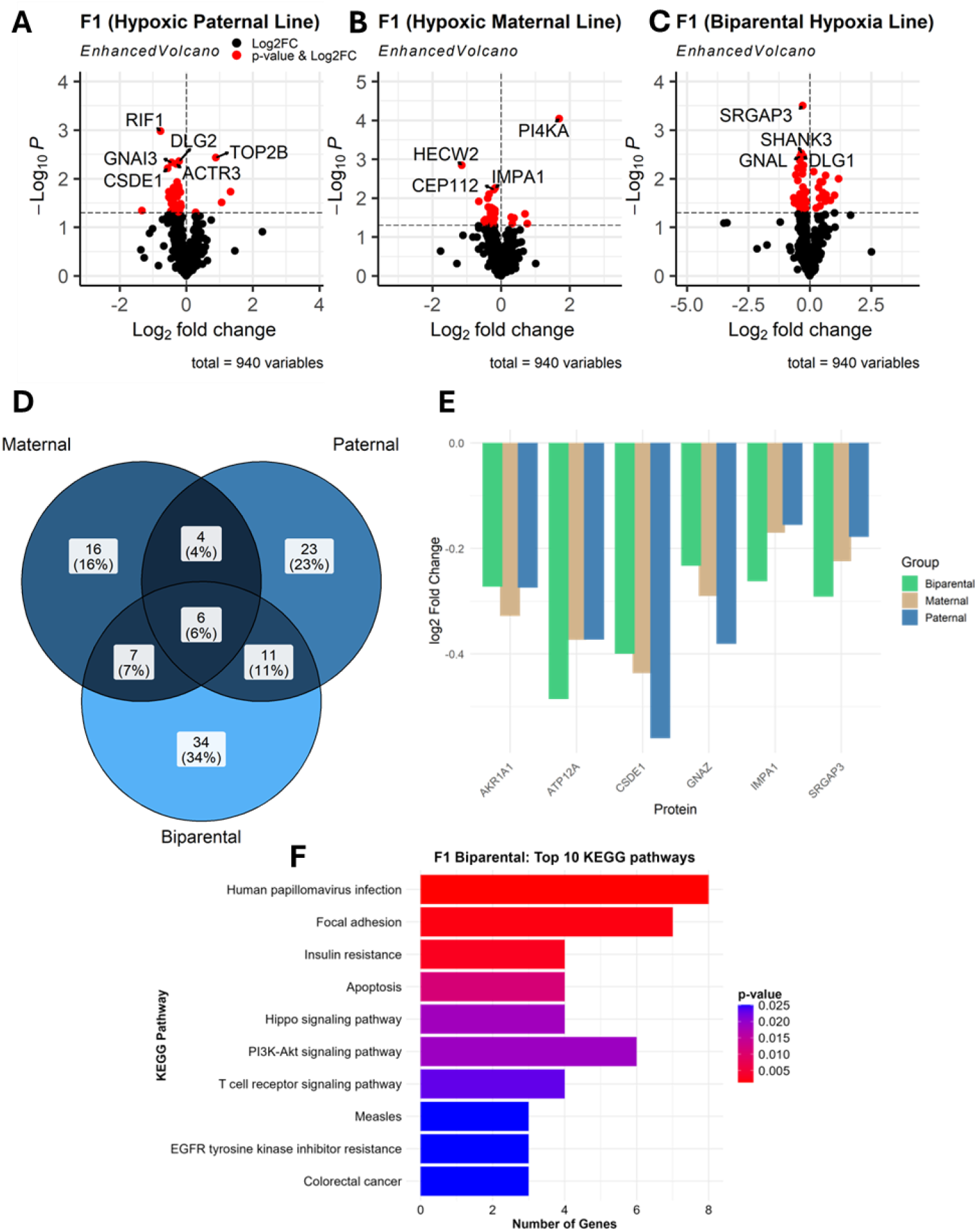
Proteomic analysis of hypoxia-induced changes in F1 mice brain cortex. **A-C)** Volcano plot showing differentially expressed (DE) proteins in paternal-hypoxic vs normoxic F1 female mice **(A)**, maternal-hypoxic vs normoxic F1 female mice **(B)**, and biparental-hypoxic vs normoxic F1 female mice **(C)**. Differential expression was defined by p-value < 0.05. **D)** Venn diagram displaying commonly DE proteins shared among paternal-, maternal-, and biparental-hypoxic F1 mice compared to normoxic controls. **E)** Bar plot illustrating the six DE proteins that are consistently altered across paternal-, maternal-, and biparental-hypoxic F1 groups relative to normoxic controls. **F**) KEGG pathway enrichment analysis showing the top 10 significantly enriched pathways for proteins differentially expressed in biparental-hypoxic vs normoxic F1 female mice.

Given its robust and consistent phenotypic protection across infarct volume and brain swelling, we focused pathway enrichment analysis on the F1 biparental group. This group also exhibited the highest number of differentially expressed proteins, increasing the power to detect enriched biological pathways.

KEGG pathway enrichment analysis in the F1 biparental group revealed significant involvement of pathways related to HPV infection, focal adhesion, insulin resistance, apoptosis, Hippo signalling, PI3K-Akt signalling, T cell receptor signalling, and EGFR signalling. Collectively, these pathways point towards alterations in cell survival, immune modulation, and intracellular signalling cascades that may underlie the observed intergenerational neuroprotection.

## 4. Discussion

Taken together, these results demonstrate that intermittent hypoxia reduces infarct volume and brain swelling in F0 mice and that this protective effect is transmitted to F1 offspring. Both maternal and paternal hypoxia exposure contributed to reduced injury, with biparental hypoxia showing the greatest protection. The observed sex-specific differences suggest distinct mechanisms of inherited resilience. These findings support the idea that intermittent hypoxia induces heritable changes that improve stroke outcomes in offspring, potentially via epigenetic mechanisms and highlighting novel avenues for stroke prevention.

Our proteomic data reveal intra-group consistency that aligns with the observed phenotypes. Normoxic F0 and F1 females clustered together, while hypoxia-exposed F1 groups (maternal, paternal, biparental) formed a distinct cluster, indicating a heritable proteomic shift. F0 females exhibited the most pronounced proteomic changes, consistent with their distinct clustering and higher number of differentially expressed proteins (DEPs), justifying their selection for pathway analysis. In F1 offspring, the biparental group demonstrated the most robust phenotypic protection and the highest number of DEPs, supporting its selection for pathway enrichment analysis. Interestingly, F0 males showed limited proteomic changes despite their protective phenotype, suggesting divergent or subtler molecular mechanisms.

Several of our findings align with existing literature, including the differential expression of *Gnai3* and *Marcksl1*, which are implicated in brain injury and ageing, respectively. Key pathways enriched in the biparental group, such as PI3K-Akt (11) and EGFR signalling (12), are established mediators in ischaemic stroke.

This study has limitations, including the exclusive focus on F1 females and the lack of temporal resolution in proteomic sampling. Nevertheless, our results suggest that hypoxic preconditioning induces sex- and generation-specific proteomic responses that may underpin intergenerational neuroprotection.

## 5. Concluding remarks

This study demonstrates that intermittent hypoxic preconditioning confers robust protection against ischaemic brain injury in both treated F0 mice and their untreated F1 offspring. The intergenerational nature of this effect was most pronounced in biparental exposure, with sex-specific differences observed in both the extent of protection and the associated proteomic responses. Proteomic profiling revealed coordinated shifts in protein expression patterns linked to ancestral hypoxia, with F0 females and F1 biparental groups showing the most prominent signatures. Enriched pathways involving metabolism, immune modulation, and intracellular signalling provide a mechanistic basis for the observed resilience.

These findings suggest that parental environmental exposures can prime offspring for improved stroke outcomes through sex- and lineage-dependent proteomic adaptations. This work adds to the growing body of evidence supporting the intergenerational impact of preconditioning and highlights potential proteins and molecular pathways for future therapeutic exploration in stroke and other neurovascular diseases.

## Author contributions

A.M. conceptualised the work. A.M.: designed the experiments; A.B.C, M.C.B., S.A., H.E.Y., A.C., carried out the animal work; F.F.I, A.B.C., M.C.B., O-N.B, D.J.B., A.A., M.D.F. and A.M analysed the data from the animal work; F.F-I, M.C.B, and E.D. analysed the proteomics dataset. F.F-I, E.K., and A.M. wrote the original draft; M.D.F., D.J.B., A.A., O-N.B reviewed and edited the manuscript. All authors have read and agreed to the published version of the manuscript.

